# Working Memory Load Modulates Neuronal Coupling

**DOI:** 10.1101/192336

**Authors:** Dimitris A. Pinotsis, Timothy J. Buschman, Earl K. Miller

## Abstract

There is a severe limitation in the number of items that can be held in working memory. However, the neurophysiological limits remain unknown. We asked whether the capacity limit might be explained by differences in neuronal coupling. We developed a theoretical model based on Predictive Coding and used it to analyze Cross Spectral Density data from the prefrontal cortex (PFC), frontal eye fields (FEF) and lateral intraparietal area (LIP). Monkeys performed a change detection task (Buschman et al., 2011). The number of objects that had to be remembered (memory load) was varied (1-3 objects in the same visual hemifield). Changes in memory load changed the connectivity in the PFC-FEF-LIP network. Feedback (top-down) coupling broke down when the number of objects exceeded cognitive capacity. Thus, impaired behavioral performance coincided with a break-down of Prediction signals. This provides new insights into the neuronal underpinnings of cognitive capacity and how coupling in a distributed working memory network is affected by memory load.

## Introduction

The number of objects that can be held in working memory (cognitive capacity) is limited (Vogel and Machizawa, 2004). Cognitive capacity is directly related to cognitive ability (Conway et al., 2003; Alloway and Alloway, 2010; Fukuda et al., 2010; Unsworth et al., 2014) and is lowered in neurological diseases and psychiatric disorders (Luck and Vogel, 2013). Therefore, studying how working memory load affects neural processing can inform our understanding of why there is a capacity limit and how cognitive function breaks down in various neurological and psychiatric diseases and disorders.

Studies of working memory load and its limits have focused on coordinated activity in frontoparietal networks known to play a major role in working memory (Klingberg et al., 2002; Todd and Marois, 2005; Palva et al., 2010; Dotson et al., 2014; Gray, 1994; Awh et al., 2006). These studies predicted capacity limits using measures of network integration (how different parts of these networks are connected together) and synchrony (Roux et l., 2012; Stevens et al., 2012). In light of recent observations that visual working memory is independent for the two visual hemifields (Buschman et al., 2011; Kornblith et al., 2016) and that changes in load have different effects on oscillatory dynamics of different frequencies (Kornblith et al., 2016), we aimed for a further understanding of working memory load on network dynamics in the frontoparietal cortex.

To that end, we re-examined LFP data from a change detection task in which working memory load was varied between one and three objects in each hemifield (Buschman et al, 2011). We previously reported (Kornblith et al.,2015) that load affected low (8-50 Hz) and high (50-100 Hz) power differently depending on time during the trial. We found a dissociation between the effects of load on lower-versus higher-frequency power and their relationship to behavior. Notably, independence between the visual hemifields was apparent in high, but not low, frequencies. Independence means that increases in stimulus load in one hemifield has no effect on the animal's ability to remember stimuli in the opposite hemifield (Buschman et al, 2011). Likewise, increasing stimulus load only effects neural activity related to stimuli in the same hemifield, not neural activity related to stimuli in the opposite hemifield (Buschman et al., 2011; Kornblith et al., 2016). Also, load effects on power were similar below and above the cognitive capacity This cannot explain abrupt decrease in behavioral performance above capacity Further, earlier power and synchrony analyses did not describe the directionality of interactions between brain areas.

Here, we aim to provide a mechanistic explanation of load effects by focusing on changes in the strength and directionality of neuronal coupling. We develop a large-scale cortical network model comprising the prefrontal cortex (PFC), frontal eye fields (FEF), and the lateral intraparietal area (LIP). This is an extension of our earlier model (Pinotsis et al., 2014; Bastos et al., 2015a) based on Predictive Coding and uses Cross Spectral Density (CSD) responses to infer changes in neuronal coupling that underlie the changes in spectral power at different frequencies. Our model addresses how load-dependent dynamics effects directed functional connectivity. It also suggests abrupt changes in neuronal coupling above capacity and a break-down of Prediction signals. Finally, it shows that functional hierarchies in large cortical networks do not necessarily change when neuronal coupling changes.

## Results

Our change detection task and behavioral results has been described in detail (Buschman et al., 2011), see also Figure 1. Two monkeys were presented with a sample array of 2 to 5 colored squares for 800 ms. This was followed by a delay period (800-to 1000-ms). After that, a test array was presented. This differed from the sample array in that one of the squares had changed colour (target). Monkeys were trained to make a saccade to the target. We analysed LFP data from the memory delay period. During this delay, there was no sensory stimulation or motor responses that might affect neuronal dynamics. We examined the relationship between dynamics and functional connectivity.

**Figure 1.**
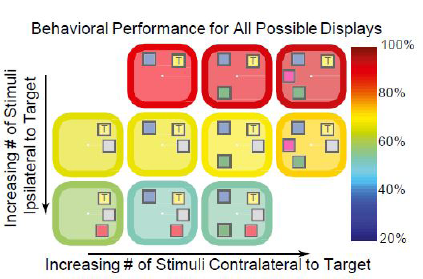
Behavioral performance (indicated by the color of the border/background) for all possible stimulus displays. Adding objects to the same side (ipsilateral) as the target (marked with a “T”) impaired performance (rows), whereas adding objects to the other side (contralateral) had no effect. This result argues for separate capacities in each hemisphere.

In our earlier work (Buschman et al., 2011; Kornblith et al., 2016), we found separate, independent capacities in the right vs left visual hemifields. Early in the memory delay, lower frequency power decreased with both ipsilateral and contralateral load but high frequency power increased only with contralateral load. By contrast, late in the memory delay, low frequency power continued to decrease with ipsilateral load but increased with contralateral load. Also, ipsilateral load effects on high frequency power were weak and there was no effect of contralateral load. To sum up, there were different effects on power in the LIP-FEF-PFC network depending on whether load was presented in the ipsilateral or contralateral visual hemifield. This motivated a separate analysis of load effects for the two hemifields. This was also supported by 1) different processing of contralateral and ipsilateral stimuli found in visual perception studies, see e.g. (Tootell et al., 1998) and 2) anatomical differences in frontoparietal connectivity for ipsilateral and contralateral connections, see e.g. (Barbas et al., 2005).). 3) Our own prior results using these same data. Behaviorally and neurophysiologically, the two hemifields seem separate and independent. Increasing stimulus load only affected behavioural performance to stimuli on the same side, not those in the opposite hemifield (Buschman et al.,2011). Neurophysiologically, we see the same thing. Stimulus information in spiking is only degraded by increases in stimulus load in the same hemifield (Buschman et al., 2011) and, likewise, effects on LFP power are hemifield dependent.

In short, we assumed that power changes found in (Kornblith et al., 2016) reflect coupling changes and analysed the effects of ipsilateral and contralateral load separately. The idea that changes in phenomenological measures, like power, reflect coupling changes is common: for example, pathological oscillations are thought to be the signature of aberrant neuronal coupling in psychiatric diseases (Uhlhaas and Singer 2012), such as autism (Dickinson et al. 2015) or schizophrenia (Gonzalez-Burgos and Lewis 2008). Further, in previous work (Miller and Cohen, 2001) we suggested that FB signals from PFC to lower areas support successful execution of working memory and decision making tasks. In this task, the animal successfully performed the task below but not above the cognitive capacity limit. We therefore expected PFC coupling to change below and above the capacity limit.

We focused on changes between 1. early vs late delay and 2. different load values (1-3 items per hemifield). We expected differences in coupling for early and late delay because our prior work using these data showed differences in power and synchrony between the early and late delay that depended on load and laterality (Kornblith et al., 2016). We therefore asked if there were any differences in PFC signals between early and late delay. We also tested different loads because performance degrades with increased load (Buschman et al., 2011).

To sum up, both electrophysiological and behavioural data suggest that coupling may change in the PFC-FEF-LIP network. The idea that changes in phenomenological measures, like the different effects on power for ipsilateral and contralateral load discussed above, reflect coupling changes is common:, for example, pathological oscillations are thought to be the signature of aberrant neuronal coupling in psychiatric diseases (Uhlhaas and Singer 2012), such as autism (Dickinson et al. 2015) or schizophrenia (Gonzalez-Burgos and Lewis 2008). Our analysis comprised three parts. First, we found the coupling *pattern* in the PFC-FEF-LIP network during the memory delay in order to determine their basic functional connectivity. Second, we asked whether the *strength* of connections changed with changes in contralateral and ipsilateral load and between the early vs late memory delay. Third, we examined *how* changes in load below vs above the animal’s behavioral capacity limit affected network connectivity.

### Functional hierarchy in the PFC-FEF-LIP network

We first examined the functional hierarchy between the PFC, FEF, and LIP. To find this hierarchy, we adapted our earlier canonical microcircuit (CMC) model (Pinotsis et al., 2014; Bastos et al., 2015a) to describe activity in the PFC-FEF-LIP network, see Figure 2.

**Figure 2.**
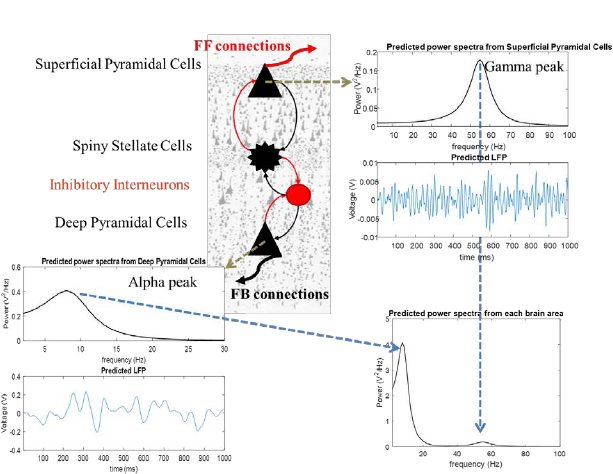
The canonical microcircuit model (CMC). The model suggests a canonical cortical architecture for the primate cortex. There are four populations of neurons (spiny stellate cells, superficial and deep pyramidal cells and inhibitory interneurons). These are connected together with excitatory (red) and inhibitory (black) intrinsic connections (thin lines). This set of populations and connections is motivated by anatomical and theoretical considerations supporting a canonical cortical microcircuitry (Douglas and Martin, 2007; Bastos et al., 2012; Pinotsis et al., 2013). Power spectra and LFPs produced by these cells are shown in the top right and bottom left plots respectively. Power spectra from each brain area are shown in the bottom right plot. Model parameters are chosen so that superficial and deep pyramidal cells oscillate at the gamma and alpha bands. These different laminar responses result from assuming different time constants (depending on GABAergic vs. glutamatergic neuromodulation) and intrinsic delay parameters for the different neuronal populations.

The CMC model is based on the Predictive Coding Model (Bastos et al., 2012) and experimental (Buffalo et al., 2011) and theoretical observations (Bauer et al., 2014; Friston et al., 2015). It builds on experimental observations that superficial and deep pyramidal cells oscillate at the gamma and alpha band respectively (Bastos et al., 2015b; Michalareas et al., 2016) and that superficial and deep pyramidal cells are the main origins of feedforward (FF) and feedback (FB) connections (Hilgetag et al., 1996; Vezoli et al., 2004). These spectral asymmetries across cortical layers, (i.e. gamma power predominant in superficial and alpha power predominant in deep layers) follows from Predictive Coding where FB connections convey Prediction signals at slower time scales (alpha) compared to bottom-up connections that convey Prediction error signals at faster time scales (gamma). Following these observations, the parameters of the CMC model were chosen so that superficial and deep pyramidal cells oscillate at the gamma and alpha band, respectively (Figure 2).

We extended the CMC model to construct a large scale model that could describe the activity in the PFC-FEF-LIP network (the “large-scale CMC model” ;Figure 3). It is an extension of the single area CMC model shown in Figure 1 and comprises FF and FB connections between PFC, FEF and LIP (red and black thick lines in Figure 3). These connections define an anatomical hierarchy (see also Experimental Procedures and Methods section). Lower areas send signal to higher areas via FF connections and receive top down input from them via FB connections. FF (respectively FB) connections are assumed to be excitatory (respectively. inhibitory). FF (respectively FB) input from area A to area B results in an increase (respectively decrease) of activity in area B that is proportional to the activity in area A. The constant of proportionality is the FF (respectively FB) coupling strength. In Predictive Coding, FF and FB signals form the basis of how the brain understands the world: according to this theory, the brain’s goal is to predict sensory inputs. Brain areas interact recurrently so that predictions (FB signals) are compared to sensory inputs and updated according to how much they deviate from them (FF signals). The theory suggests that this iterative process is repeated until deviations are minimised. Thus, FF (sensory) input excites higher cortical areas. FB signals inhibit FF inputs and allow only FF signals that were not predicted to be passed forward.

**Figure 3.**
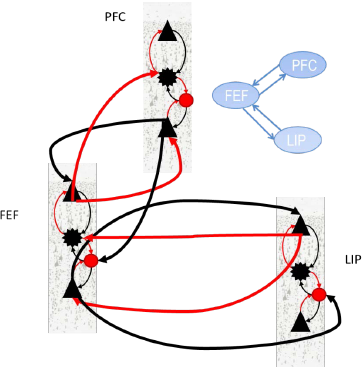
Large scale CMC model. The model describes the large-scale structure of the primate cortex. Different brain areas are connected with extrinsic connections (thick lines). Intrinsic connections are as described in Figure 2. Extrinsic connections are as follows: Feedforward (FF) connections are assumed to originate in superficial layers and target input spiny stellate cells and deep pyramidal cells. Feedback (FB) connections are assumed to originate from deep layers and target superficial pyramidal cells and inhibitory interneurons. This pattern of extrinsic and intrinsic coupling has been shown to explain activity in parietal and frontal areas (Heinzle et al., 2007; Ma et al., 2012; Phillips et al., 2015; Ranlund et al., 2016; Díez et al., 2017). We have omitted extrinsic connections between PFC and LIP in the Figure and depicted one out of four possible connection patterns that corresponds to results from anatomical studies (Hilgetag et al., 2016). This is variant “FEF" of the large-scale CMC model (see main text for details).

To sum so far, our large-scale CMC model predicts oscillatory interactions and hierarchical relations in the PFC-FEF-LIP network based on FF and FB coupling between brain areas and local oscillatory dynamics within each area. It also assumes that FF and FB signals propagate at different frequencies and convey Prediction Errors and Predictions respectively Thus, it allows us to test whether changes in phenomenological measures, like power, reflect coupling changes. It also allows us to link coupling changes due to memory load to Predictive Coding. Increasing FB (resp. FF) signals with increasing load would correspond to stronger Predictions (resp. Prediction errors). In this context, failure to perform the task when the number of stimuli exceeds cognitive capacity, implies a failure in Prediction signals. Thus, we expected FB from PFC to break down above the capacity limit.

The anatomical hierarchy shown in Figure 3 follows recent studies that exploit differential laminar source and termination patterns and tract tracing experiments to obtain the hierarchical distribution of brain areas (Hilgetag et al., 2016; Markov et al., 2014; Medalla and Barbas, 2006). However, whether the functional hierarchy will follow the anatomical hierarchy is not clear. Functional hierarchies are not as robust as anatomical hierarchies and are often task-dependent (Buschman and Miller, 2007; Bastos et al., 2015b).

To find the functional hierarchy in the PFC-FEF-LIP network, we fitted the large-scale CMC model to CSD data from trials with the same memory load. This data contained information about oscillatory interactions in different frequency bands (Kornblith et al., 2016). For model fitting, we used Dynamic Causal Modeling (DCM; (David et al., 2006; Pinotsis and Friston, 2014; Moran et al., 2015; Pinotsis et al., 2016; Garrido et al., 2009; Kiebel et al., 2009). DCM is a standard approach for model fitting. It has been widely used to determine the directionality of information flow and functional hierarchy in brain networks (Gluth et al., 2015; Hare et al., 2011; Hillebrandt et al., 2014; Li et al., 2014; Smith et al., 2006). Specifically, DCM has been applied to the analysis of neuronal activity in frontal and parietal areas and during functions ranging from attention to memory, decision making and psychiatric diseases, similarly to the frontoparietal network and working memory task considered here, see (Mechelli et al., 2004; Garrido et al., 2009; Wang et al., 2010; Jacques et al., 2011; Vossel et al., 2012; FitzGerald et al., 2015).

To find the functional hierarchy, we used Bayesian model comparison (BMC, see Friston et al., 2007). BMC is a process comprising 1. model fitting and 2. computation of model evidence. Model evidence is a mathematical quantity that expresses how likely each a model is for a given dataset. Usually one considers a set of models (model space) and finds the model with highest evidence. We first fitted different variants of the large-scale CMC model (Figure 3) to our data. These model variants differed in the connections between PFC, FEF and LIP. They are shown in Figure 4A and describe all possible functional hierarchies. They are called “ALL”, “FEF”, “LIP” and “PFC” respectively. Model “FEF” is the model where FEF is connected to PFC and LIP and there are no direct connections between PFC and LIP. This was the coupling of the model shown in Figure 3 and is what one would expect from anatomical studies (Hilgetag et al., 2016). These alternatives are described by the other three models. These are similar to model” FEF”, where FEF is replaced by PFC and LIP. Model "ALL" assumed that all areas were connected and information flows in FF and FB directions between all areas. When two areas are connected with both FF and FB connections, we say that they are connected with reciprocal (R) connections.

**Figure 4.**
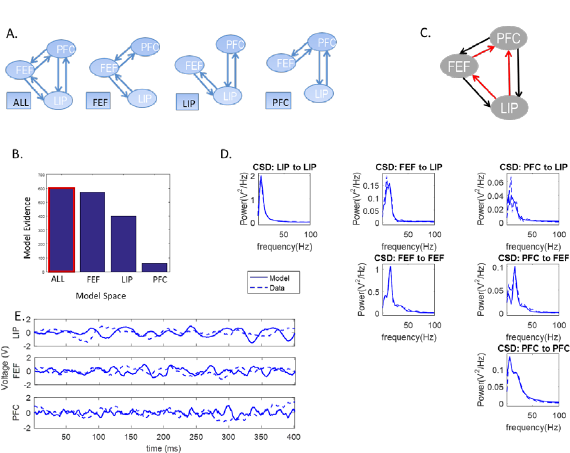
A. Possible functional hierarchies in the PFC-FEF-LIP network. B. Bayesian model comparison (BMC) results after fitting variants of the large-scale CMC model to trials with one contralateral object during the early delay period. C. The model with highest evidence was model “ALL”. All brain areas occupy the same hierarchical level. D. Model fits to CSD data. E. Using posterior parameter estimates, we simulated LFPs. In all plots, dashed lines depict model predictions and solid lines depict observed data (CSD or LFPs).

To find the model evidence, BMC uses an approximation called Free Energy. Conceptually, BMC can be thought of as a generalization of classical model comparison approaches, e.g. Bayesian Information Criterion (BIC). The difference is in the cost function used. BMC uses Free Energy that also includes a complexity term in addition to an accuracy term. For a model to have the highest evidence both terms should be maximized: the accuracy term is maximized when the model fits the data best (i.e. it has the smallest error). The complexity term is maximized when all model parameters are necessary for fitting the data. If a model has parameters that are not necessary, this term will not be maximum and therefore the evidence for that model will be lower. The reason is that unnecessary parameters will have large posterior correlations between them. Each parameter does not explain the data in a unique way (similarly to coefficients of determination in classical statistics, posterior correlations quantify the explanatory power of model parameters in Bayesian statistics). These correlations will enter into the complexity term and make it smaller (for more details see Friston et al., 2007). Even if the model with the highest number of parameters fits the data best (has the maximum accuracy) this model will not have the highest evidence if some parameters are unnecessary (the complexity term will not be maximum).

We performed BMC between variants of the large-scale CMC model. This allowed us to find the model that best fit the data and whose parameters were necessary for fitting the data (i.e. the model that did not “over-fit” the data). We fitted the four models of Figure 4A to four different datasets. The first two datasets included CSD data from trials with one contralateral object, from the early (500-900ms after sample onset) or late (1100-1500ms after sample onset) delay period. The last two datasets included CSD data from trials with one ipsilateral object (and the same delay periods). The winning model had the highest evidence among all models considered. In general, the difference in model evidence between (winning) model A and its runner up B is useful because it immediately yields the probability that model A is more likely than model B (this is called exceedance probability of model A vs B) ^1^. It can be shown mathematically, that if this difference is bigger than 3, the exceedance probability is equal to 1, that is the winning model is 100% more likely than its runner up and any other model that was considered. A summary of the fitting process is included in Section “Dynamic Causal Modeling” of Supplementary Material. This process has also been described in detail in several earlier publications, see e.g. (Friston et al., 2012; Pinotsis et al., 2014).

We first fitted CSD data from trials with one contralateral object during the early delay period (different memory loads are presented below). The results of our analysis are shown in Figure 4. Model fits are shown in Figure 4D. Plots show alpha and gamma power model fits: in most cases model predictions (solid lines) fully overlapped with experimental data (dashed lines). This is not surprising as priors have been carefully chosen to accurately reproduce alpha and gamma activity (Bastos et al., 2012; Friston et al., 2015). Small discrepancies between data and model fits occurred only when CSD power was weak (∼ 0.05V^2^/Hz, top right panel in Figure 4D). Model fitting yielded posterior parameter estimates. Including these estimates in our model, we obtained simulated LFPs. These are shown with solid lines in Figure 4E. Observed LFPs are shown with dashed lines.

Figure 4B shows the model evidence for the four models tested (corresponding to possible hierarchies shown in Figure 4A). The winning model was model “ALL”: all areas were connected with reciprocal connections (highlighted with a red frame in Figure 4B, shown in Figure 4C). Model fits and simulated LFPs show a good fit to experimentally observed data (Figures 4D and 4E, respectively).

Supplementary Figures 1–3 show the corresponding results for contralateral load and late delay and ipsilateral load and early and late delay respectively. These are very similar to Figure 4. Although load effects on power were different between early and late delay and for contralateral and ipsilateral load, see Figure 5 in (Kornblith et al., 2016), we found that the functional hierarchy did not change between the early and late delay periods and was also the same for contralateral and ipsilateral objects. Model fits and LFPs are also shown in Supplementary Figures 1B–3B and 1C–3C and are very similar to Figures 4D and 4E.

**Figure 5.**
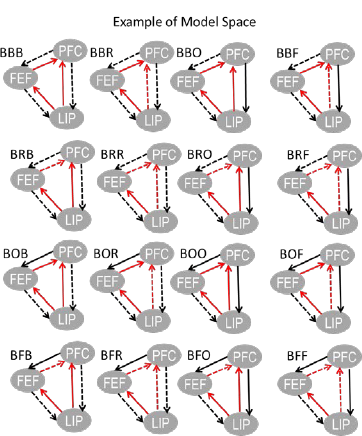
Plots showing variants of model “ALL”. These correspond to the 16 variants included in first two lines of Table 1. Each variant had an acronym. The letters in the acronym correspond to connections that were allowed to change for different load conditions. These are also shown with dashed lines. Solid lines depict connections that were not allowed to change.

For a contralateral object, the difference in model evidence between model “ALL” and its runner up was ΔF *=* 29 for early delay and ΔF = 111 for late delay (Figure 4B and Supplementary Figure 1A). For ipsilateral object, the difference in model evidence of the winning model “ALL” with respect to the runner up was ΔF = 60 during early delay and ΔF = 21 during late delay (Supplementary Figures 2A and 3A). Because ΔF was bigger than three, model “ALL” had exceedance probability equal to 1 for all four datasets considered. These results were robust to using trials with different memory loads (see Supplementary Figure 4). In all cases, model “ALL” had the highest evidence. This means that the functional hierarchy in the PFC-FEF-LIP network did not change when changing memory load and for different parts of the delay period despite the different load effects on spectral power^2^.

To sum so far, we found that all three brain areas in the PFC-FEF-LIP network had reciprocal functional connections. In other words, all three areas were on the same hierarchical level. The *pattern* of FF and FB connections (functional hierarchy) did not change with memory load and for early vs late delay. Next, we compared alternative variants of the winning model of the first part of our analysis (model “ALL”) where we allowed a different subset of FF and FB connections to change with load (and the rest of the connections were left unaffected). This revealed changes in the strength of functional connections with changes in memory load^3^.

### Feedforward and feedback coupling strengths changed load and time

Above, we saw that the model “ALL” best captured the functional hierarchy between PFC, FEF, and LIP (i.e., they all had reciprocal, R, connections with each other). Here, we test whether FF, FB, or R coupling in the early vs late memory delay was affected by working memory load. We did so using BMC to compare variants of model “ALL”. In these model variants, different FF, FB or R connections were allowed to change for different load conditions. These model variants are described by an acronym shown in the entries of Table 1. They corresponded to all possible connections that could change with increasing object load and included models where connections did not change. There were 64 such variants. The first 16 variants (first two lines of Table 1) are also depicted in Figure 5. The same variant was fitted to CSD data for all contralateral and ipsilateral load conditions and from data from early or late in the memory delay. Coupling parameters were allowed to change progressively between the lowest and highest load conditions. Coupling changes between different load conditions were assumed to be linear increments (increases or decreases) to coupling corresponding to lowest load. In other words, load changes were assumed to have modulatory effects on cortical coupling and are called trial specific effects in DCM. This is similar to trial specific effects in fMRI literature (Coderre and van Heuven, 2013; Den Ouden et al., 2008; Gordon et al., 2015).

**Table 1^4^.**

|  |  |  |  |  |  |  |  |
| --- | --- | --- | --- | --- | --- | --- | --- |
| 1:BBB | 2: BBR | 3: BBO | 4: BBF | 5:BRB | 6: BRR | 7: BRO | 8: BRF |
| 9:BOB | 10: BOR | 11: BOO | 12: BOF | 13:BFB | 14: BFR | 15: BFO | 16: BFF |
| 17:RBB | 18: RBR | 19: RBO | 20: RBF | 21:RRB | 22: RRR | 23: RRO | 24: RRF |
| 25:ROB | 26: ROR | 27: ROO | 28: ROF | 29:RFB | 30: RFR | 31: RFO | 32: RFF |
| 33:OBB | 34: OBR | 35: OBO | 36: OBF | 37:ORB | 38: ORR | 39: ORO | 40: ORF |
| 41:OOB | 42: OOR | 43: OOO | 44: OOF | 45:OFB | 46: OFR | 47: OFO | 48: OFF |
| 49:FBB | 50: FBR | 51: FBO | 52: FBF | 53:FRB | 54: FRR | 55: FRO | 56: FRF |
| 57:FOB | 58: FOR | 59: FOO | 60: FOF | 61:FFB | 62: FFR | 63: FFO | 64: FFF |

Each variant had an acronym, please see Table 1. For example, in variant BBF, the feedback connections between LIP and FEF and FEF and PFC and the feedforward connections between LIP and PFC were allowed to change with load. Connections that were not allowed to change with increasing memory load were depicted with solid lines. Connections that were allowed to change were depicted with dashed lines. Using BMC, we identified the most likely model (that is, the set of connections affected by contralateral and ipsilateral load and in the early vs late delay) among the 64 alternatives (see Figure 6A for contralateral load and early delay and Supplementary Figures 5A–7A for contralateral load and late delay and ipsilateral load and early and late delay respectively). These Figures include bar plots of model evidence. We call the space of all possible variants “model space”. This is shown in the horizontal axis and is the same as in Table 1. Figures 6B and 6C and Supplementary Figures 5B–7B and 5C–7C show model fits to CSD data and simulated and observed LFPs respectively. These are similar to results in Figure 4D and 4E and Supplementary Figures 1B–3B and 1C–3C.

**Figure 6.**
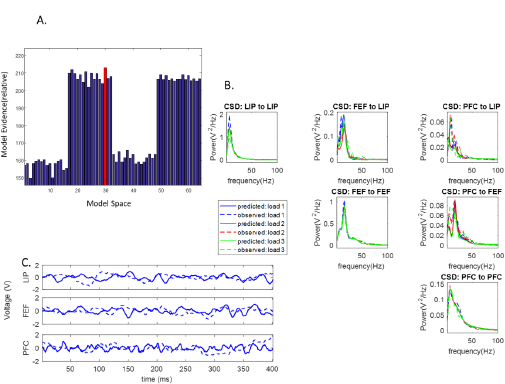
Contralateral WM Load Effects on FF and FB coupling in the PFC-FEF-LIP network during early delay. Plots follow the format of Figure 3. A. Bayesian model comparison (BMC) results after fitting the 64 variants of model “ALL” included in Table 1. B. model with highest evidence was model “RFR”. D. Model fits to CSD data. C. Simulated and observed LFPs. In all plots, dashed lines depict model predictions and solid lines depict observed data (CSD or LFPs).

The winning models are shown in Figure 7A(i)–(iv) and Supplementary Figures 5B–7B: RFR and ROB were the winning models for contralateral load during early and late delay and BFB and RRR were the winning models for ipsilateral load during early and late delay. Model evidence of winning models is shown with red bars. The difference in model evidence between them and their runner ups was Δ*F* = 1 and Δ*F* = 24 for contralateral load during early and late delay and Δ*F* = 0.7 and Δ*F* = 4 for ipsilateral load respectively. The corresponding model exceedance probabilities were 74% and 100% for variants RFR and ROB and 65% and 100% for variants BFB and RRR respectively.

**Figure 7.**
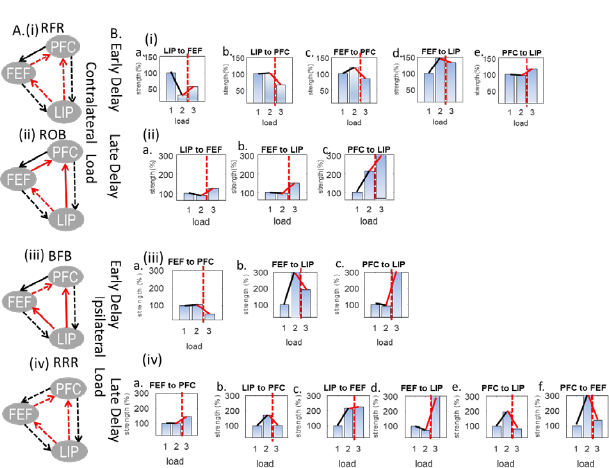
A. Models that had the highest evidence for each experimental condition. Dashed lines denote connections that were modulated by load. RFR and ROB were the winning models for contralateral load during (i) early and (ii) late delay and BFB and RRR were the winning models for ipsilateral load during (iii) early and (iv) late delay. B. Changes in neuronal coupling strengths due to changes in (i) contralateral load during early delay; (ii) contralateral load during late delay; (iii) ipsilateral load during early delay; (iv) ipsilateral load during late delay. Coupling strengths corresponding to the lowest load are shown with blue bars. The cognitive capacity limit is shown with a vertical red dashed line. Strength changes below (resp. above) the capacity limit are shown with black (resp. red) lines.

What these models showed was that changing contralateral load affected R coupling between LIP and FEF and FB coupling between PFC and LIP throughout the delay period (Figure 7A(i) and 7A(ii)). However, FF input to PFC from the other two brain areas was affected by contralateral load only during early delay (Figure 7A(i)). Also, changing ipsilateral load affected FB coupling between LIP and the other two areas and FF coupling between FEF and PFC throughout the delay period (Supplementary Figures 7A(iii) and 7A(iv)). During late delay, on the other hand, all connections in the network were affected by changing load: on top of the above connections, FF coupling between LIP and the other two areas and FB coupling between FEF and PFC changed with ipsilateral load (Supplementary Figure 7A(iv)).^5^

To sum this analysis, we identified different sets of connections that were affected by increasing memory contralateral and ipsilateral load during different parts of the delay period. Having established load-specific changes in connection strengths, we can now proceed to our last analysis. In this last set of analyses we examine in greater detail how coupling changed with changes in load below and above the animal’s working memory capacity.

### Increases and decreases in feedforward and feedback coupling strengths below and above capacity

Here, we explore the changes in coupling between areas as a result of changes in load. To organize this discussion, we distinguish changes below (from load 1 to load 2) and above (from load 2 to load 3) the animal’s behavioral capacity. There were as many coupling parameters in our model as thick lines in winning model “ALL”, see Figure 4C.

The results below were obtained from the same model fits as in the previous section. We used model fits of the winning model only (winning models are shown in Figure 7A). In the previous section, the winning model (and the other 63 alternatives of Table 1) was fitted to CSD data for all load conditions simultaneously. Coupling strengths were allowed to change progressively between the lowest and highest contralateral and ipsilateral load. We also used CSD data obtained during early and late delay. The set of coupling strengths that changed with load determined the winning model in each case. Below, we discuss these strengths and their progressive changes, see Figure 7B.

Coupling strengths corresponding to the lowest load (one contralateral or ipsilateral object) were rescaled so that they were equal to 100%. The three blue bars in each panel show coupling strength for load 1, 2 and 3 (from left to right). This reveals progressive changes in the same coupling strength as load increases. Strengths were normalized with respect to the lowest load condition. Besides the blue bars, each panel includes three additional lines connecting adjacent bars: black (shows coupling changes below the capacity limit), solid red (shows coupling change above the capacity limit) and dashed red (separates coupling below and above the capacity limit). Recall that FB connections were inhibitory while FF connections were excitatory (these are shown with black and red arrows connecting brain areas in Figure 4C). In Figures 7B(i) (resp. 7B(ii)), we show changes in coupling strengths during the early (resp. late) delay period when contralateral load changes. In Figures 7B(iii) (resp. 7B(iv)), we show the corresponding changes when ipsilateral load changes.

Almost all parameters (15/17) were significantly modulated above the capacity limit: they showed marked increases or decreases of at least 25% or more. We will call these changes “strong” as opposed to other changes that we will call “weak”. Below the capacity limit fewer parameters were strongly modulated (9/17). We first focused on changes in coupling strength below the capacity limit. These changes can be readily seen by focusing on the slope of black lines in Figures 7B(i) and 7B(ii): almost horizontal (resp. oblique) lines correspond to weak (resp. strong) changes. Changes in connections involving PFC followed a consistent spatiotemporal pattern regardless of whether load change was contralateral or ipsilateral: they were weak during early delay and strong during late delay signaling bigger PFC involvement closer to the decision time (i.e., during late delay)^6^. Conversely, changes in the other FF and FB connections (between LIP and FEF) followed the opposite pattern: they were strong during early delay and weak during late delay but for contralateral load only (for ipsilateral load they were strong throughout the delay period).

Specifically, for contralateral load during early delay, we observed strong changes in the FF and FB connections between LIP and FEF but not PFC, compare black lines in Figures 7B(i)a and 7B(i)d with black lines in Figures 7B(i)b, 7B(i)c and 7B(i)e. Similarly, for ipsilateral load, compare black line in Figure 7B(iii)b with black lines in Figures 7B(iii)a and 7B(iii)c. This also means that excitatory input to PFC did not increase due to higher load (black lines in Figures 7B(i)b, 7B(i)c and 7B(iii)a). At the same time, FEF input from LIP decreased with increasing load (black line in Figure 7B(i)a) and input from FEF to LIP increased (black lines in Figures 7B(i)d and 7B(iii)b).

For contralateral load during late delay, the above pattern of connection changes was reversed: connections between LIP and FEF showed weak modulations with load (black lines in Figures 7B(ii)a and 7B(ii)b). At the same time, FB input from PFC to LIP showed a strong increase with increasing load (black line in Figure 7B(ii)c). Similarly, for ipsilateral load FF and FB PFC connections also showed a strong increase with load (black lines in Figures 7B(iv)b, 7B(iv)e and 7B(iv)f). However, in contrast to what we observed for contralateral load, connections between LIP and FEF continued to show strong modulations (as in the early delay period, see black lines in Figures 7B(iv)c and 7B(iv)d).

Interestingly, above the cognitive capacity limit (when going from load two to load three), changes in connections in the PFC-FEF-LIP network were strong. The only exception was the LIP-FEF connections during early delay for contralateral load and late delay for ipsilateral (red lines in Figures 7B(i)d and 7B(iv)c). We will see below that connections between LIP and FEF showed the opposite pattern of changes above capacity in comparison to their pattern below capacity. Also, signals to and from PFC were affected by load during both early and late delay (below capacity they were affected by load only during late delay). Most importantly, FB signals from PFC and FEF were modulated differently for contralateral and ipsilateral load.

During early delay, FF input to PFC from the other two brain areas was strongly reduced above the capacity limit, see red lines in Figures 7B(i)b, 7B(i)c and 7B(iii)a. Similarly, FB input from PFC to LIP increased above capacity during early delay (red lines in Figures 7B(i)e and 7B(iii)c). This was also the case for contralateral load during late delay (red line in Figure 7B(ii) c). However, for *ipsilateral* objects and late delay FB signals from PFC broke down: they showed a strong *reduction* (as opposed to increase in all other cases) when exceeding the capacity limit (red lines in Figures 7B(iv)e and 7B(iv)f). This was accompanied by a strong reduction (break down) in FF input from LIP to PFC (red line in Figure 7B(iv)b). These were the only cases where coupling above the capacity limit was very similar to coupling for lowest load. FF input from FEF on the other hand showed an increase (red line in Figure 7B(iv)a). FF input from LIP to FEF also increased above capacity regardless of object hemifield and delay period (red lines in Figures 7B(i)a, 7B(ii)a and 7B(iv)c). FB input from FEF reduced for early (red lines in Figures 7B(i)d, 7B(iii)b) and increased for late delay above capacity for both contralateral and ipsilateral objects(red lines in Figures 7B(ii)b, 7B(iv)d).

Above we described coupling changes with increasing load. To quantify how likely these changes were we used posterior probabilities (coupling estimates were found using DCM which is a Bayesian approach for model fitting). These are shown in Supplementary Figures 8 and 9. They are the probabilities of a significant non-zero change with respect to the coupling strength for the lowest load. Because all parameters were normalized to the lowest load condition, we included posterior probabilities only for Load 2 and Load 3. These are shown in matrix form. Columns correspond to the brain areas from which connections originated and rows to areas where they terminated. The number of these matrices is one less than the number of possible loads (there are no probabilities for the lowest load). The posterior probability of changes in coupling strengths for contralateral load ranged between 53-100% (resp. 62-100%) for early (resp. late) delay, see Supplementary Figure 8. The posterior probabilities for coupling parameters for ipsilateral load ranged between 71-100% (resp. 54-100%) for early (resp. late) delay, see Supplementary Figure 9.

## Discussion

We studied the effects of changing working memory load on neuronal dynamics during a change detection task. We analysed CSD data obtained using LFPs from frontal and parietal areas, namely PFC, FEF and LIP. Activity in this frontoparietal network has been found to consistently change with training (Goldman-Rakic, 1995; Li et al., 1999) and has been associated with cognitive capacity (Rottschy et al., 2012).

We followed up on earlier work (Buschman et al., 2011; Kornblith et al., 2016), where we had found that neuronal activity in high, but not low, frequencies reflects independent processing of ipsilateral and contralateral objects and changes substantially between early and late delay period. Independent processing of objects in different hemifields has also been confirmed by (Matsushima and Tanaka, 2014) and is supported by early anatomical studies (Goldman-Rakic and Schwartz, 1982). In (Kornblith et al., 2016), neuronal activity changes with load were captured as spectral power effects. However, these effects were similar below and above the cognitive capacity, which appears at odd with a reduction in behavioral performance observed when capacity is exceeded. Further, earlier power and synchrony analyses did not describe the directionality of neuronal interactions. Here, we aimed at a mechanistic explanation of load effects by focusing on changes in the strength and directionality of neuronal coupling. We extended our earlier model based on Predictive Coding (CMC model; Pinotsis et al., 2014; Bastos et al., 2015a) and used it to analyze Cross Spectral Density data. The CMC model can predict oscillatory interactions and hierarchical relations in the PFC-FEF-LIP network based on FF and FB coupling between brain areas and local oscillatory dynamics within each area. It has been validated pharmacologically (Muthukumaraswamy et al., 2015), using data from single-gene mutation channelopathy (Gilbert et al., 2016) and aging studies (Cooray et al., 2014a; Moran et al., 2014). The model has also explained the manipulation of sensory expectation and attention engaging frontoparietal networks in healthy subjects and patients (Auksztulewicz and Friston, 2015; Cooray et al., 2014b; Díez et al., 2017; Phillips et al., 2015; Ranlund et al., 2016). A very similar model was recently used to explain context ‐dependent dynamics in hierarchical brain networks (Mejias et al., 2016).

We first studied the basic functional hierarchy in the PFC-FEF-LIP network. We determined its form and asked whether this changed with memory load and time during the delay period. Anatomical connections provide the substrate for functional connections but functional hierarchies can be different than anatomical hierarchies. They can be task-dependent and even change during a task (Buschman and Miller, 2007; Bastos et al., 2015b) as a result of goal-directed behaviour (Miller, 1999; Miller and Cohen, 2001) and of processing abstract information (Koechlin et al., 2003). Also, there are reciprocal anatomical connections between frontal areas are other frontal and parietal areas (Medalla and Barbas, 2006; Hilgetag et al., 2016). Some studies have placed PFC at the top and parietal areas at the bottom of functional hierarchies in visual perception tasks (Bastos et al., 2015b; Michalareas et al., 2016). However, the functional hierarchy in the change detection task we studied here was unknown.

To find the functional hierarchy, we compared variants of our model corresponding to different hierarchical relations between PFC, FEF and LIP using Bayesian model comparison (BMC, Friston et al., 2007). We found that PFC, FEF and LIP had reciprocal functional connections (they were at the same hierarchical level). This result was the same regardless of memory load and time during the delay period. However, load effects on power were different of low for contralateral and ipsilateral objects and early vs. late delay (Kornblith et al., 2016). Therefore, it might well be that although the functional hierarchy remained the same across trials with different load and throughout the delay period, the *amount* of signal transmitted through FF and FB connections, that is, the *strength* of FF and FB connections, changed with load and time.

Thus, we then identified different subsets of FF and FB connections whose strength changed with load during different parts of the delay period. We used BMC to compare different models corresponding to all possible combinations of connections that might be affected by load. After finding the most likely model, we focused on the corresponding changes in coupling strengths. These explain the weak load effects on power (1-2% power change per added object) found in (Kornblith et al., 2016) without changing the functional hierarchy.

We found that below the capacity limit connections involving PFC were affected later than connections involving other frontal and parietal areas for contralateral and ipsilateral load. During early delay, connections between LIP and FEF were strongly affected by load while connections involving PFC did not change much. FF input from LIP decreased with increasing load while FB input to LIP increased. This could be related to the fact that receptive fields observed in LIP are unilateral and have a narrow spatial tuning (Platt and Glimcher, 1998). During late delay, connections involving PFC were strongly modulated for contralateral and ipsilateral load but connections between LIP and FEF were not affected for contralateral load. However, when ipsilateral load changed, changes in connections between LIP and FEF remained strong during late delay (similarly to early delay). This could be related to the widespread and more dense patterns of ipsilateral as opposed to contralateral connections to frontal areas (Barbas et al., 2005). Based on the above results, our model predicts that, below the capacity limit, PFC engages strongly in network activity only close to the decision time (above capacity, PFC engages throughout the delay period, see below). Further, as load increased, we observed increases in both FF input to PFC and FB signals from PFC to other frontal and parietal areas. These reflected increased FF drive due to higher load and increasing FB stabilizing signals from PFC to counteract increased in cognitive demands (load) due to increased FF drive in earlier areas. They are similar to earlier modeling results (Macoveanu et al., 2006; Edin et al., 2009; Wei et al., 2012).

Above the cognitive capacity limit, connections that we had previously identified to be affected by load changes showed strong modulations. Connections involving PFC were affected by load throughout the delay period. Importantly, FB connections were modulated differently by contralateral and ipsilateral load. FB stabilizing signals from PFC increased above capacity for contralateral load but were significantly reduced (broke down) for ipsilateral load. This could explain reduced behavioral performance when the total number of objects in the same (but not the opposite) hemifield as the target object exceeded the capacity limit found by (Buschman et al., 2011). This also provides an interesting link to Predictive Coding. Break down of Prediction signals coincided with impaired behavioral performance. This difference in coupling changes while changing contralateral vs ipsilateral load supports earlier findings about independent capacities of the two hemifields (Buffalo et al., 2011; Matsushima and Tanaka, 2014). Stabilizing signals from FEF to LIP also broke down above capacity for ipsilateral, but not contralateral load. This supports an important role of FB from frontal areas in successful performance. Interestingly, FB signals from FEF broke down earlier than PFC FB signals (these broke down closer to decision time). This might be related to the fact that loss of information about object identity in PFC occurs later than other frontal areas, see (Buschman et al., 2011).

To sum up, we found that neuronal coupling changes as a result of changing the number of objects maintained in working memory These changes are dynamic and evolve as the time for behavioral response (decision) approaches. We also found that FB coupling breaks down when the number of ipsilateral objects is above the cognitive capacity limit and that this occurs first in parietal and then frontal areas. These results shed new light in coupling changes that might underlie reduced cognitive capacity and behavioral performance. They also suggest network-specific pathological changes in neuronal coupling that might occur in various neurological and psychiatric diseases and disorders (Luck and Vogel, 2013).

## Acknowledgements

This work was supported by NIMH R37MH087027, NIMHR01MH091174 and the Picower Institute Innovation Fund.

## Supplementary Material

**Supplementary Figure 1:**
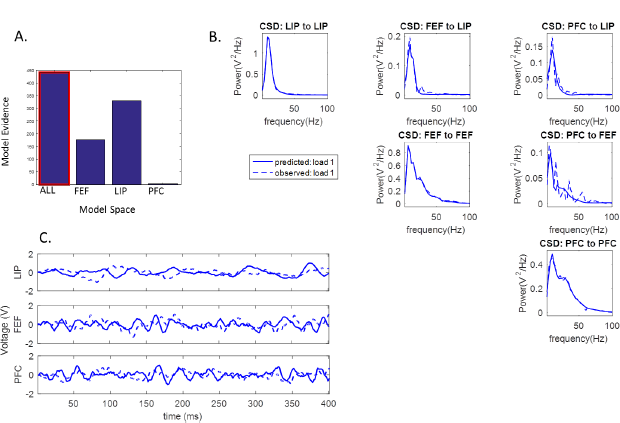
Functional hierarchy and Model Fits for contralateral Load and Late Delay. A. Bayesian model comparison results after fitting variants of the large-scale CMC model to trials with one contralateral object during the late delay period. B. Model fits to CSD data. C. Simulated and observed LFPs. In all plots, dashed lines depict model predictions and solid lines depict observed data.

**Supplementary Figure 2:**
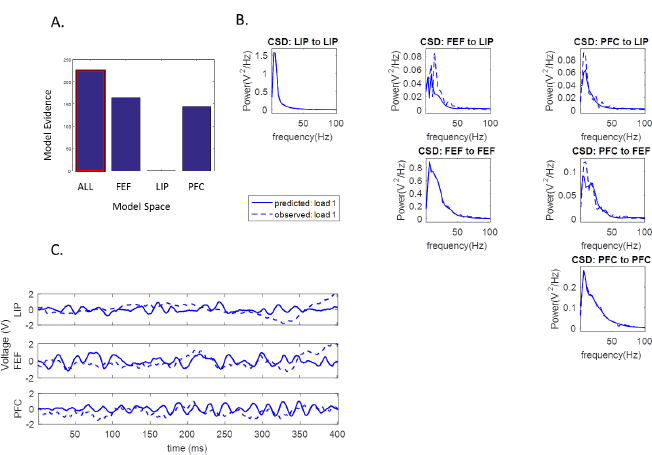
Functional hierarchy and Model Fits for Ipsilateral Load and Early Delay. A. Bayesian model comparison results after fitting variants of the large-scale CMC model to trials with one ipsilateral object during the early delay period. B. Model fits to CSD data. C. Simulated and observed LFPs. In all plots, dashed lines depict model predictions and solid lines depict observed data.

**Supplementary Figure 3:**
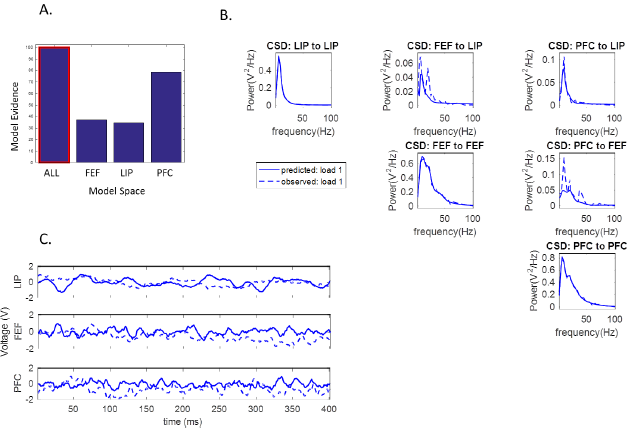
Functional hierarchy and Model Fits for Ipsilateral Load and Late Delay. A. Bayesian model comparison results after fitting variants of the large-scale CMC model to trials with one ipsilateral object during the late delay period. B. Model fits to CSD data. C. Simulated and observed LFPs. In all plots, dashed lines depict model predictions and solid lines depict observed data.

**Supplementary Figure 4:**
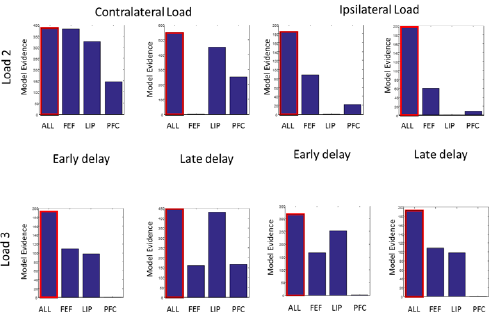
Bayesian model comparison for finding functional hierarchy. Bayesian model comparison (BMC) results after fitting variants of the large-scale CMC model to trials with two (upper panels) or three (lower panels) contralateral (left panels) and ipsilateral (right panels) objects for different parts of the delay period.

**Supplementary Figure 5:**
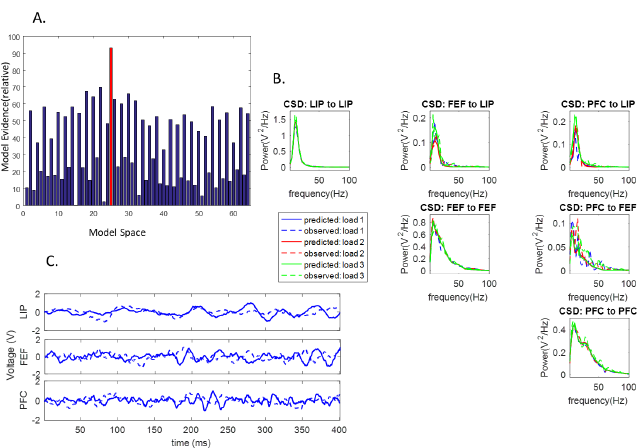
Contralateral WM Load Effects on FF and FB coupling in the PFC-FEF-LIP network during late delay. Plots follow the format of Figure 3. A. Bayesian model comparison (BMC) results after fitting the 64 variants of model “ALL” included in Table 1. B. Model fits to CSD data. E. Simulated and observed LFPs. In all plots, dashed lines depict model predictions and solid lines depict observed data.

**Supplementary Figure 6:**
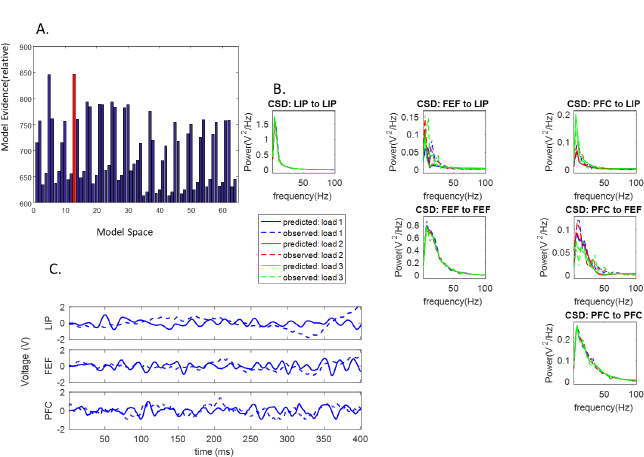
Ipsilateral WM Load Effects on FF and FB coupling in the PFC-FEF-LIP network during early delay. Plots follow the format of Figure 3. A. Bayesian model comparison (BMC) results after fitting the 64 variants of model “ALL” included in Table 1. B. Model fits to CSD data. E. Simulated and observed LFPs. In all plots, dashed lines depict model predictions and solid lines depict observed data.

**Supplementary Figure 7:**
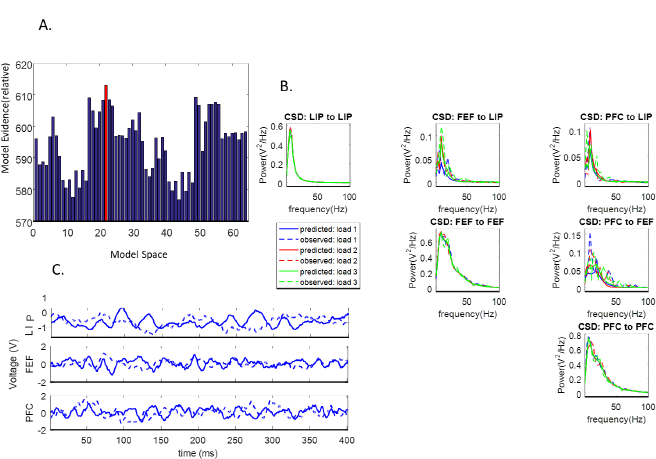
Ipsilateral WM Load Effects on FF and FB coupling in the PFC-FEF-LIP network during early delay. Plots follow the format of Figure 3. A. Bayesian model comparison (BMC) results after fitting the 64 variants of model “ALL” included in Table 1. B. Model fits to CSD data. C. Simulated and observed LFPs. In all plots, dashed lines depict model predictions and solid lines depict observed data.

**Supplementary Figure 8:**
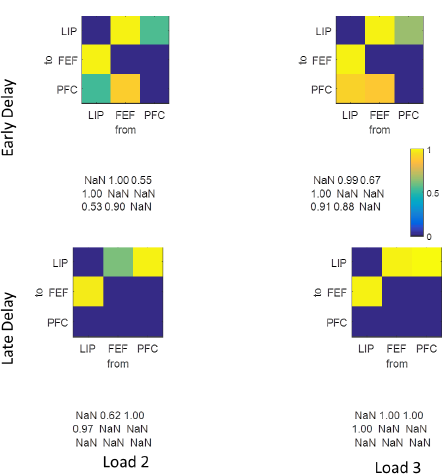
Posterior Probability of Changes in Coupling With Contralateral Load. Posterior probability of changes in neuronal coupling with contralateral load. Upper (respectively lower) panels show estimates for early (respectively late) delay. Left (respectively right) panels show estimates for two (respectively three) objects.

**Supplementary Figure 9:**
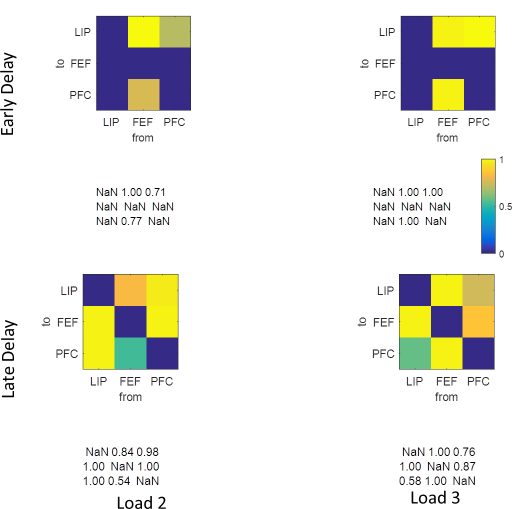
Posterior Probability of Changes in Coupling With Ipsilateral Load. Posterior probability of changes in neuronal coupling with ipsilateral load. Upper (respectively lower) panels show estimates for early (respectively late) delay. Left (respectively right) panels show estimates for two (respectively three) objects.

**Supplementary Figure 10:**
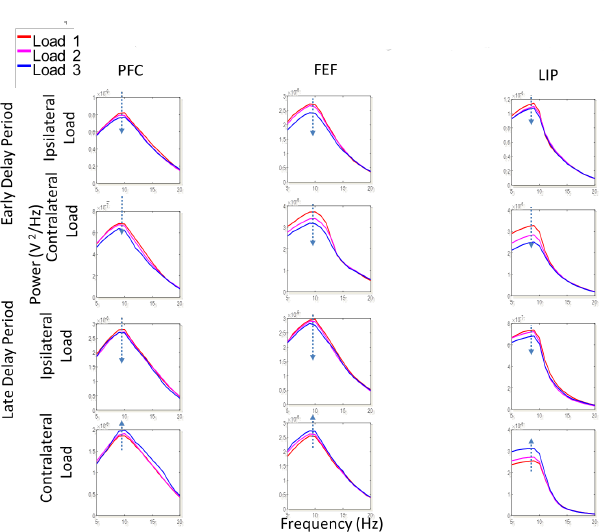
Power Spectra of Significantly Modulated Recorded Sites. Power spectra of significantly modulated sites. Red, pink, blue and yellow curves correspond to power spectra for working memory load 0,1,2 and 3 respectively. The first (resp. last) two rows correspond to the early (resp. late) delay period. First, second and third column show power spectra obtained from PFC, FEF and LIP respectively. For illustration purposes, power spectra are shown for a range between 5 and 20 Hz where differences in power with load were evident. It should be noted, however, that for all our analyses we used the whole spectrum 2-100Hz and fitted our model to CSD data from that spectrum.

**Supplementary Figure 11:**
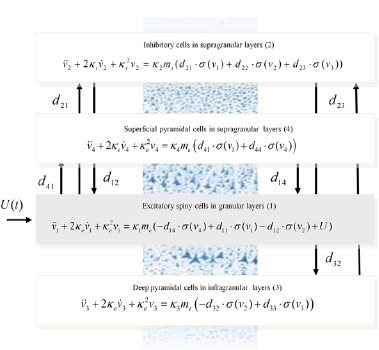
Equations of the CMC model. This figure shows the evolution equations that specify a Canonical Microcircuit (CMC) neural mass model of a single source. This model contains four populations occupying different cortical layers: there are two pyramidal cell populations that are thought to be the sources of forward and backward connections in cortical hierarchies (see also main text). Second-order differential equations mediate a linear convolution of presynaptic activity to produce postsynaptic depolarization. This depolarization gives rise to firing rates within each sub-population that provide inputs to other populations.

## Supplementary Experimental Procedures and Methods

### Summary

Two adult male monkeys (monkey Sp, 13 kg; monkey Si, 6kg) were handled in accordance with National Institutes of Health guidelines and the Massachusetts Institute of Technology Committee on Animal Care. They were trained to perform a change detection task. This task required the monkeys to fixate a central dot. After this fixation period (500ms), a sample array of 2–5 colored squares was presented for 800ms. After a memory delay that ranged from 800ms to 1000ms, the test array appeared: this differed from the sample array in that one of the squares in the sample array had changed color in the test array at random. This was called the target square. The monkey was trained to make a single, direct saccade to the target square.

Six new object locations were chosen each day. Square locations were chosen from 6 positions (3 per hemifield) in any single session in each visual hemifield, ranging from ±75 angular degrees from the horizontal meridian and between 4 and 6 degrees of visual angle (dva) from fixation. Objects were colored squares 1 dva on a side. Two colors were chosen for each location every day. Given behavioral evidence for the independence of working memory representations in each hemifield, in each trial, we manipulated the number of squares in each hemifield between 1 and 3. One of these squares was the target and the rest were the distractors. Contralateral (ipsilateral) load was defined as the total number of squares in the opposite (same) hemifield as the target. Both the location and colors were changed each day, to prevent monkeys from adopting any long-term memorization strategy. The colors were drawn from a predefined population of 14 colors in a random manner as long as 2 colors were not too difficult to discriminate at a particular location (i.e., red and pink were never paired). This process ensured a large degree of variety in the objects used on a particular day and thus required the animals to encode and hold in memory the array presented on each trial and detect its change rather than memorizing fixed object response associations. The location of the target object was chosen randomly on each trial. Early in training the number of total objects in the array was chosen randomly. We equalized the number of trials in each of the experimental condition (one, two or three objects in each hemifield). Therefore, the constellation of the objects used was pseudorandomly chosen such that both the distribution of trials with one, two, or three objects in the target’s hemifield and the distribution of total number of objects in a trial were flattened. This process did not alter the probability of the target location. Both monkeys performed the task well above chance and with similar accuracy. Correct trials were those that the monkey made a saccade to the target square. Correct trials were used in this paper.

### Neurophysiological Recordings and Data

Neuronal signals were recorded simultaneously across a maximum of 50 electrodes distributed across frontal and parietal cortex. LFP data were obtained from electrodes in prefrontal cortex (PFC), frontal eye fields (FEF) and the lateral intraparietal areas (LIP). The signal from each electrode was transformed to a local field potential (LFP) by filtering between 3.3 and 88 Hz and down sampling to 1 kHz. LFPs were referenced to ground. Power-line artefacts at 60, 120, and 180 Hz were estimated and subtracted from the data. We computed CSD data (2-100 Hz) from LFP using FieldTrip’s^7^ wavelet transform method. We analyzed data from two different periods in the delay, 500-900ms and 1100-1500ms after sample offset. These periods showed stationary activity associated with object maintenance. To identify the specific electrodes that were used in the coupling analysis, we first selected a subset of electrodes residing in PFC, FEF or LIP according to anatomy and neurophysiological responses (Buschman et al., 2011). To identify LIP neurophysiologically, we trained the animals on a delayed saccade task. Electrodes were considered to be within LIP only if a neuron isolated from that electrode showed memory delay activity during that task. Microstimulation was used to demarcate FEF from PFC (Bruce and Goldberg, 1985). Only sites that had thresholds of stimulation amplitudes <50 μA were classified as belonging to FEF. Anterior sites within the frontal well were classified as belonging to PFC. In general, stimulation at PFC sites did not elicit eye movements even at the highest current amplitude tested (150 μA). Within each subset of electrodes, we calculated the contrast-to-noise ratio (CNR) during the early and late delay periods for each electrode (Cui et al., 2011). This CNR was calculated from the pooled signal-to-noise ratio corresponding to experimental conditions with different working memory loads *(i=2,3)*

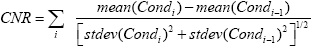

The CNR calculates the signal amplitude change between experimental conditions during the periods of interest, divided by the pooled standard deviation. Larger CNR indicates that signal change between conditions is larger. In each brain area, we chose a single electrode from each brain area based on two criteria: (i) It had the highest CNR; (ii) CSD power changed strictly monotonically between conditions in accord with our earlier results (Kornblith et al., 2016). It should be noted that carefully selecting electrodes did not bias the coupling analysis below. Statistical significance of results is quantified in terms of relative model evidence (Bayesian analogue of odds ratio) and parameter posterior probability. This is the cumulative probability of the conditional distribution obtained after model fitting.

In (Kornblith et al., 2016) we performed a detailed time frequency analysis using a generalised linear model and wavelet transforms across all electrodes, time points and frequencies and found strictly monotonic power changes with memory load in each brain area. Changes were significant but small (between 1% to 7%). For the current analysis, we selected electrodes that exhibited the same monotonic relationships as those found in (Kornblith et al., 2016), see Supplementary Figure 10: red, pink, blue and yellow curves correspond to power spectra for working memory load 0,1,2 and 3 respectively. The first (resp. last) two rows correspond to the early (resp. late) delay period. First, second and third column show power spectra obtained from PFC, FEF and LIP respectively. For illustration purposes, power spectra are shown for a range between 5 and 20 Hz where differences in power with load were evident. It should be noted, however, that for all our analyses we used the whole spectrum 2-100Hz and fitted our model to CSD data from that spectrum. Computation of CSD data proceeds by averaging over time points and trials. This means that not all power changes identified using time frequency analysis in (Kornblith et al., 2016) will be evident in the CSD data. These relationships were only evident at the single electrode level: taking averages of CSD data over all electrodes in a brain area did not show monotonic changes.

### Dynamic Causal Modeling

Dynamic Causal Modeling(DCM) is a framework for the analysis of neuroimaging data (Moran et al., 2013). It allows one to infer neuronal mechanisms and architectures by combining ideas from dynamical systems with Bayesian inference (Friston et al., 2003; Pinotsis et al., 2012). We here used DCM to a neural mass model to the CSD data reported in (Kornblith et al., 2016) known as the Canonical Microcircuit model (CMC), see Figure 1. We used routines developed earlier by us and colleagues that are freely available as part of SPM toolbox^8^. We extended this model to describe neural activity in the PFC-FEF-LIP network, see Figure 2. The differential equations appearing in this model are included in Supplementary Figure 11. Model parameters include gains on each channel’s time constant and coupling parameters. For a detailed explanation of model parameters we refer the reader to (Pinotsis et al., 2013, 2014).

Given this set of differential equations we applied a linear stability analysis in the spectral domain to obtain CSD data predicted by the CMC model. We followed the approach of (Kornblith et al., 2015), where instead of power, we used coupling parameters to characterize changes in neuronal activity due to changing load. Note that (Kornblith et al., 2015) found power changes of about 1-2% due to changing load and did not find different effects between the two monkeys. Based on this result and the robustness of our model, we did not distinguish between the two monkeys. This allowed us to establish a mapping from coupling parameters to CSD data (Friston et al., 2012; Pinotsis and Friston, 2014). This had the form of a probabilistic model that we then fitted to measured data; here CSD data from the PFC-FEF-LIP network. We thus obtained posterior estimates densities of coupling parameters that allowed us to describe working memory load effects on neuronal dynamics. To obtain these estimates we used a variational Bayesian scheme that maximizes the negative Free energy, see (Friston et al., 2007). This scheme performs a coordinate descent on the negative Free energy. It minimizes it with respect to a variational density q(e). After convergence, the variational density approximates the true posterior *q*(eI *y*, *m*) ∝ *p*(y|e, *m*)*p*(el*m*) and the Free energy approximates the model evidence *F* = −ln *p*(*y|m*). Thus, the variational density provides posterior estimates of coupling parameters and Free energy can be used for model comparison.

## Footnotes

1 To obtain the exceedance probability from the difference in model evidence one has to apply a sigmoid function, see (Kass and Raftery, 1995).

2 A careful reader might question if finding model “ALL” (the model with most parameters) as the winning model might be the result of overfitting. In the main text, we laid out technical arguments about how the particular cost function (Free Energy) used for model comparison prevents this. We also noted that we obtained the same result using 12 different datasets (Figure 4B and Suppl. Figures 1A–3A and 4). On top of these arguments, we note that we found a different model as the winning model using the same datasets but changing the threshold of high pass filtering. During our preliminary investigations (not shown), we had found winning model “FEF” by using trials where ipsilateral load was varied and focusing on low frequency responses only (2-50Hz).

3 Note that in the analysis above we fitted the large-scale CMC model to data from trials with the same memory load. Our goal was to test whether certain connections were present or not. In the analysis below, we fitted the model to data from trials with different memory load simultaneously. This allowed us to focus on changes of model parameters with increasing load.

4 Each variant had an acronym. The first letter in this acronym corresponds to the connections that were allowed to change with load between LIP and FEF. The second letter corresponds to the connections that were allowed to change between FEF and PFC. The third letter corresponds to the connections that were allowed to change between PFC and LIP. The letters F, B and R correspond to feedforward, feedback and reciprocal connections respectively. The letter O corresponds to connections that were not allowed to change.

5 The results of Figure 6 and Supplementary Figure 7 show that models with R of FF connections between LIP and FEF are favoured in the corresponding model comparisons. These are the models 17-32 and 49-64, see also rows 3-4 and 7-8 in Table 1. These results can also be useful for family-wise inference which we will consider in future work.

6 This general trend had one exception: the FF connection from FEF to PFC which was weakly modulated when increasing ipsilateral load during late delay, see Figure 6(iv)a.

7 http://www.ru.nl/neuroimaging/fieldtrip/

8 http://www.fil.ion.ucl.ac.uk/spm/software/spm12/

